# Intercontinental comparison of ferns community assemblages in Malaysian and Nigerian tropical forests

**DOI:** 10.1101/2021.08.04.455095

**Authors:** Gbenga F. Akomolafe, Rusly Binti Rosazlina, Zakaria Rahmad, Fatai A. Oloyede

**Affiliations:** School of Biological Sciences, Universiti Sains Malaysia, 11800 Penang, Malaysia; Department of Plant Science and Biotechnology, Federal University of Lafia, Nigeria; Department of Botany, Obafemi Awolowo University, Ile-Ife, Nigeria

**Keywords:** ecological zones, ferns diversity, intercontinental, species richness, tropical forests

## Abstract

Research on ferns ecology has gained attention in the last decade, yet there is paucity of information on the intercontinental comparison of ferns community across two continents. This study focused on comparing the ferns community assemblages in tropical forests of Malaysia and Nigeria, thereby assessing the patterns of the species richness and diversity across the two continents. The diversity and taxonomic compositions of ferns were assessed using 180 plots of size 10 m x 10 m in each country. The species richness and other diversity indices were determined using the combined forests data for each country and for the individual forests. The observed and rarefied–extrapolated fern species richness are significantly higher in Malaysian forests than Nigerian forests. Also, the other diversity indices (Simpson index, Margalef index, and Fisher’s alpha) are significantly higher in Malaysian forests except Shannon index which showed no significant difference between the two biogeographic regions. There is a very low similarity in the taxonomic composition of ferns between the two biogeographic areas, although the similarity in composition increased with increasing taxonomic levels (genus and family levels). Terrestrial and epiphytic ferns are more dominant than the other life forms in the two countries. Since the two countries receive varying degrees of environmental factors, we then hypothesize that these observed differences are due to climatic differences as well as historical and evolutionary processes.

## Introduction

Pteridophytes (Ferns) are known to be an essential part of the biodiversity and vegetations of tropical forest ecosystems [1]. They originated from the old world tropics and have colonized other regions of the world [2]. Ferns generally constitute the substantial biomass of many tropical and subtropical forests of the world [3]. Their occurrence and abundance in these tropical regions are largely dependent on moisture availability [4]. Apart from their biodiversity roles in the ecosystems, they are also useful to mankind in diverse ways such as for ornamentals, medicines, and food [5, 6]. Over the years, there have been several collections of ferns across many tropical forests of Malaysia which have led to the documentation of over 1165 fern species out of the 4400 species reported for South East Asia [7, 8]. Similarly, ferns are known to be found widely across many ecological zones of Nigeria, constituting about 165 species, 64 genera, and 27 families [9]. Although several factors such as unplanned urbanization, farming, and mineral exploitations have threatened their existence in many parts of Nigeria [10].

Malaysia and Nigeria are both located within the tropics, yet they both differ in some climatic conditions. For instance, researches have reported that tropical forests in Africa receive lesser amount of rainfall than the ones of Southeast Asia [11, 12]. In the same vein, the rainfall extends through the year in Southeast Asian tropical forests as compared with that of West Africa forests. This results in a higher relative humidity in Asian tropical forests than in African tropical forests [13]. These climatic differences have caused variations in the floristic composition between the two regions. There is little information on the ferns differences between Asian and African forests as researchers are more focused on trees, lianas and other plant types [14].

Due to the evolutionary and climatic differences of the two continents, single research that assesses floristic data from these two tropical regions can boost our understanding of biogeographic patterns in ferns ecology. Single researches that cover more than one continent are regarded as more effective in drawing the intercontinental patterns of species ecology than individual researches that only focus on one specific continent [14]. Hence, our study focused on determining the intercontinental patterns in ferns community assemblages between Malaysian and Nigerian tropical forests. Consequently, the following questions were asked: (1) does ferns diversity and community structure differs between Malaysian and Nigerian forests? (2) Are there any similarity in the ferns taxonomic composition between Malaysian and Nigerian forests?

## Materials and Methods

### Study Area

The study was conducted in some tropical recreational forests and University campuses in Nigeria and Malaysia. In Nigeria, the ferns species were collected from the campus of the Obafemi Awolowo University (OAU), Ile-Ife, Osun State and the reserved forest near Ikogosi warm spring, Ekiti State (Fig. 1). This University campus is located in Ile-Ife which lies between latitude 7°32′N and longitude 4°32′E. The campus was originally sited on a primary forest which has now been transformed into secondary forest due to the level of human encroachments [15]. Ile-Ife city falls within the tropical rain forest zone of Nigeria which receives an average annual rainfall of 1400 to 1500 mm and a mean annual temperature ranging between 27°C to 34°C. The tropical rain forest vegetation is characterized mainly by abundant trees with few woody shrubs. Ile-Ife like other cities in Nigeria has two main seasons including the dry and rainy seasons. The city experiences rainy season between March and October whereas it experiences dry season between November and early March [16]. The second site, Ikogosi warm spring forest is located at Ikogosi, Ekiti State. It has a geographical boundary of latitude 7°34’ N and longitude 4°58’E. This warm spring has been recognized as a recreational center which houses both primary and secondary forests. One major interesting thing about the warm spring is the confluence point between it and another cold spring.

**Figure 1:**
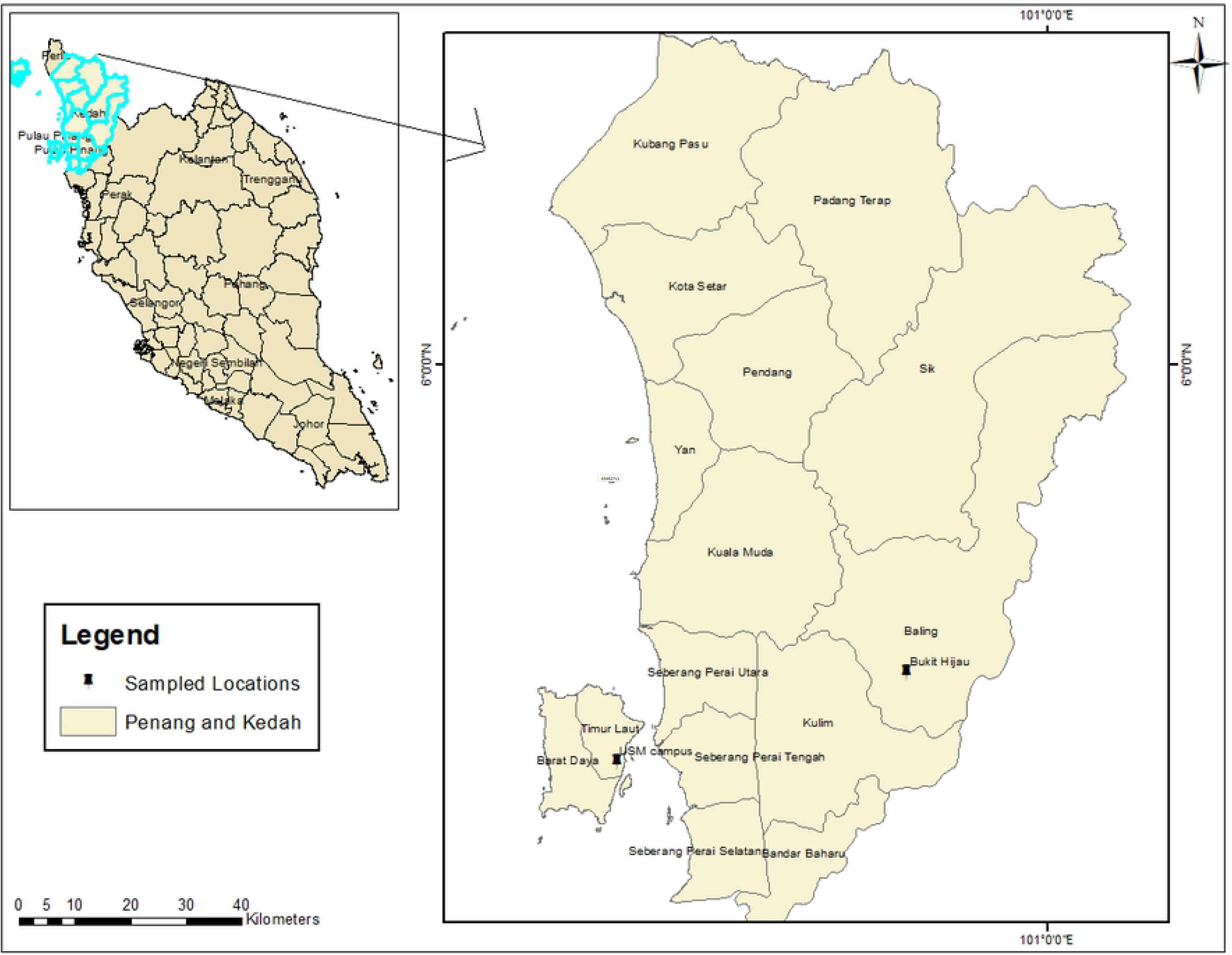
Study area map of Peninsular Malaysia showing USM campus and Bukit Hijau forests.

In Malaysia, the ferns were collected from the main campus of the Universiti Sains Malaysia (USM), Penang Island and Bukit Hijau recreational forest, Kedah (Fig. 2). Both sites are located within the Peninsular Malaysia. This peninsular is known to be the floristically richest part of Indomalesian sub-kingdom [17]. This campus is situated on about 252.7 hectares of land. The vegetation comprises extensively large canopy trees. This city has a tropical climate with an average annual rainfall of almost 2670 mm [18]. Also, it receives a daily temperature range of 24°C to 32°C and relative humidity of 70 to 90 %. The second site which is Bukit Hijau recreational forest is located at 42 km away from Baling town, Kedah, Peninsular Malaysia. It has a geographic boundary of latitude 5°30^Ꞌ^N and longitude 100°46^Ꞌ^E). This forest was established as a recreational forest in 1959, and it has been used for activities such as picnics, campsites, eco-tourism, and educational activities [7]. The forest has also housed wildlife, such as elephants, tigers, tapirs, monkeys, deer, birds, and squirrels [19]. The forest is characterized by lowland and slightly hill dipterocarp vegetation with a waterfall and is elevated at 150–300 m above sea level.

**Figure 2:**
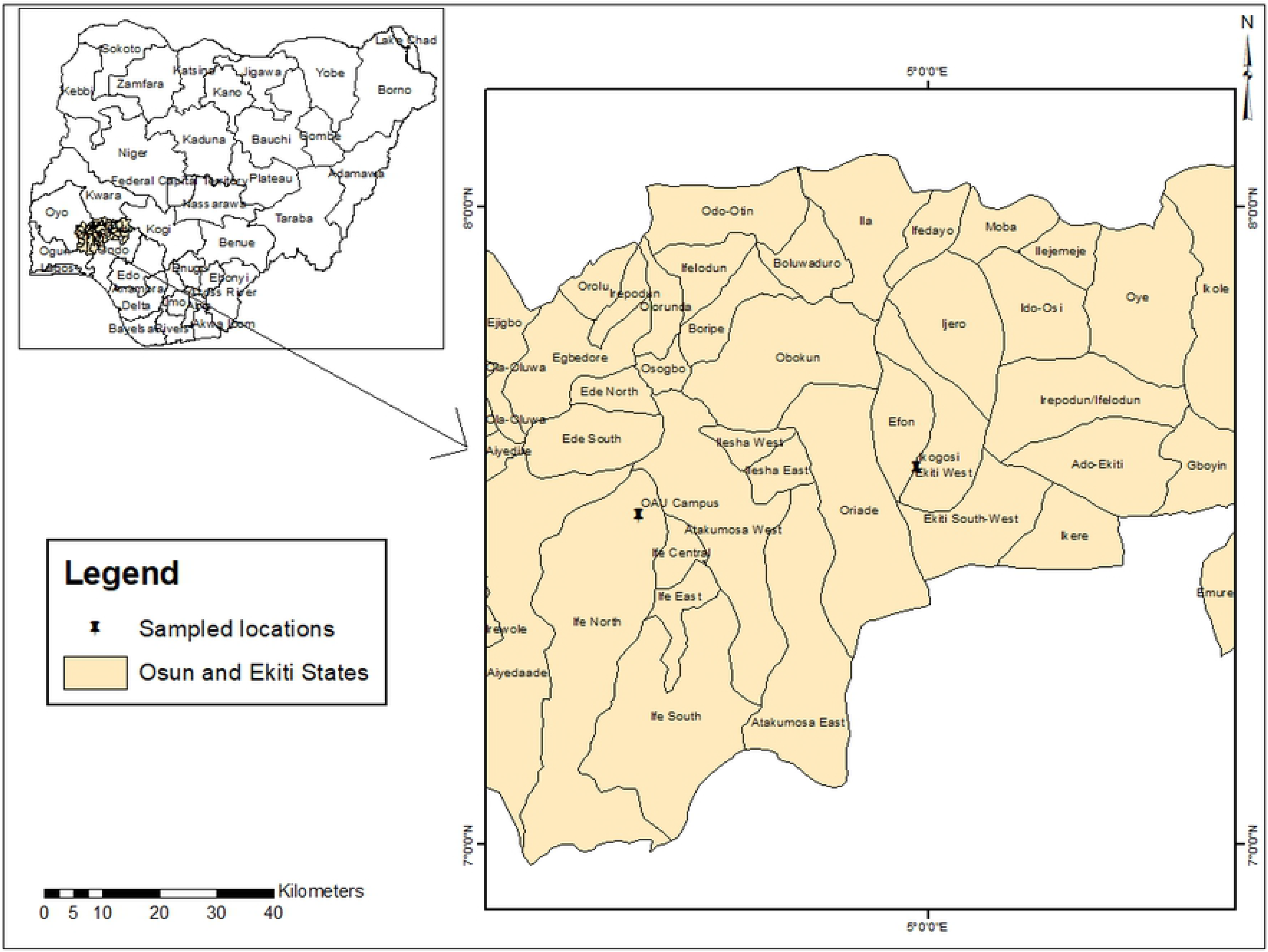
Study area map of Nigeria showing OAU campus and Ikogosi warm spring forest.

### Ferns Sampling Technique

In each site, ferns were sampled at three different areas within the forested areas. This includes the undisturbed area, less-disturbed area and most-disturbed area. These areas were selected based on the observation of the rate of human encroachments and infrastructural developments. The sampling was designed to cover all the three areas. Thirty plots of size 10 m x 10 m were established in each area, giving rise to a total of 90 plots per site and 180 plots in each country. A preferential non-random method of sampling which ensured that at least one individual fern was captured in each plot, was adopted for the study [9].

In all the established plots, the abundance of individual ferns found were noted and categorized as terrestrial, aquatic, epiphytes and lithophytes. Some of the ferns were identified directly on the field. Those with difficult identification were pressed and identified at the herbaria of Universiti Sains Malaysia and Federal University of Lafia. The fern species were thereafter identified using taxonomic flora [20, 21, 22] and online database of International Plant Names Index. The voucher specimens were deposited in the respective herbarium depending on the country for references. The conservation status of each identified fern was assessed from the redlist database of the International Union for the Conservation of Nature (IUCN).

### Data Analyses

The relative frequency of each fern species was calculated for the two biogeographic areas. In order to have a considerable comparison of the species richness between the two campuses, a non□asymptotic rarefaction-extrapolation analysis which is a species richness evaluator was employed [14]. This was done using the individual-based abundance data for each fern species. Significant difference in the rarefied-extrapolated fern species richness between the countries was determined by the confidence intervals of the species accumulation curves, constructed using 50 bootstrap replicates. This analysis was achieved with the aid of iNEXT software (online version) [23]. An overlap in the confidence intervals of the curves indicates the absence of statistical significance difference between the species richness of the two countries. Significance difference in the species richness is only established when the confidence intervals of the curves are non-overlapping [7]. This analysis is very important as it eradicates bias which is usually encountered in field sampling. The ferns diversity indices such as Shannon index, Simpson index and Margalef index were quantified for each country using paleontological statistics (PAST) 3.0 software. Pairwise permutation tests were used to determine the significant differences in the diversity indices between the countries.

Sorensen’s similarity index was used to determine the intercontinental similarity in the ferns compositions at the three taxonomic levels (species, genus and family) between the two studied areas. The life forms of the ferns were associated with the two countries using a non-metric multidimensional scaling (nMDS) analysis with Bray-Cutis dissimilarity measures with the aid of the PRIMER 7 software. Variations in the distribution of the ferns species across the two continents were determined using the principal component analysis (PCA) of the PAST software.

## Results and Discussion

A total of 54 ferns species were observed in Malaysian forests while total of 27 ferns were observed in Nigerian forests (Table 1). This difference in the ferns species is significant as revealed by the pairwise permutation test (*P* = 0.001). In the same vein, the rarefied-extrapolated species richness shows that the Malaysian forests are richer in ferns than Nigerian forests (Fig. 3a). The rarefied and extrapolated species richness curves for both countries reached asymptote showing that there was an adequate sampling of the ferns in the forests. This is also evident by the equal values of the observed and rarefied-extrapolated species richness of ferns of both countries. All the diversity indices of the combined forests of Malaysia are significantly higher than those of Nigerian forests (pairwise permutation test: *P* = 0.001) except the Shannon diversity index which showed no significant difference (*P ˃* 0.05). These trends generally indicated that Malaysian forests are richer and more diverse in fern species than the Nigerian ones. This could be due to the significant roles moisture and other microclimatic conditions play in the occurrence and distribution of ferns in tropical forests [4]. South East Asian countries particularly Malaysia is known to receive more rainfall throughout the year than West African ones [14, 13]. This may account for the differences in the richness of ferns between the two continents. It is also important to state that the two countries are more diverse in fern species as ecosystems with Shannon index of 2 and above are considered more diverse [24].

**Figure 3a:**
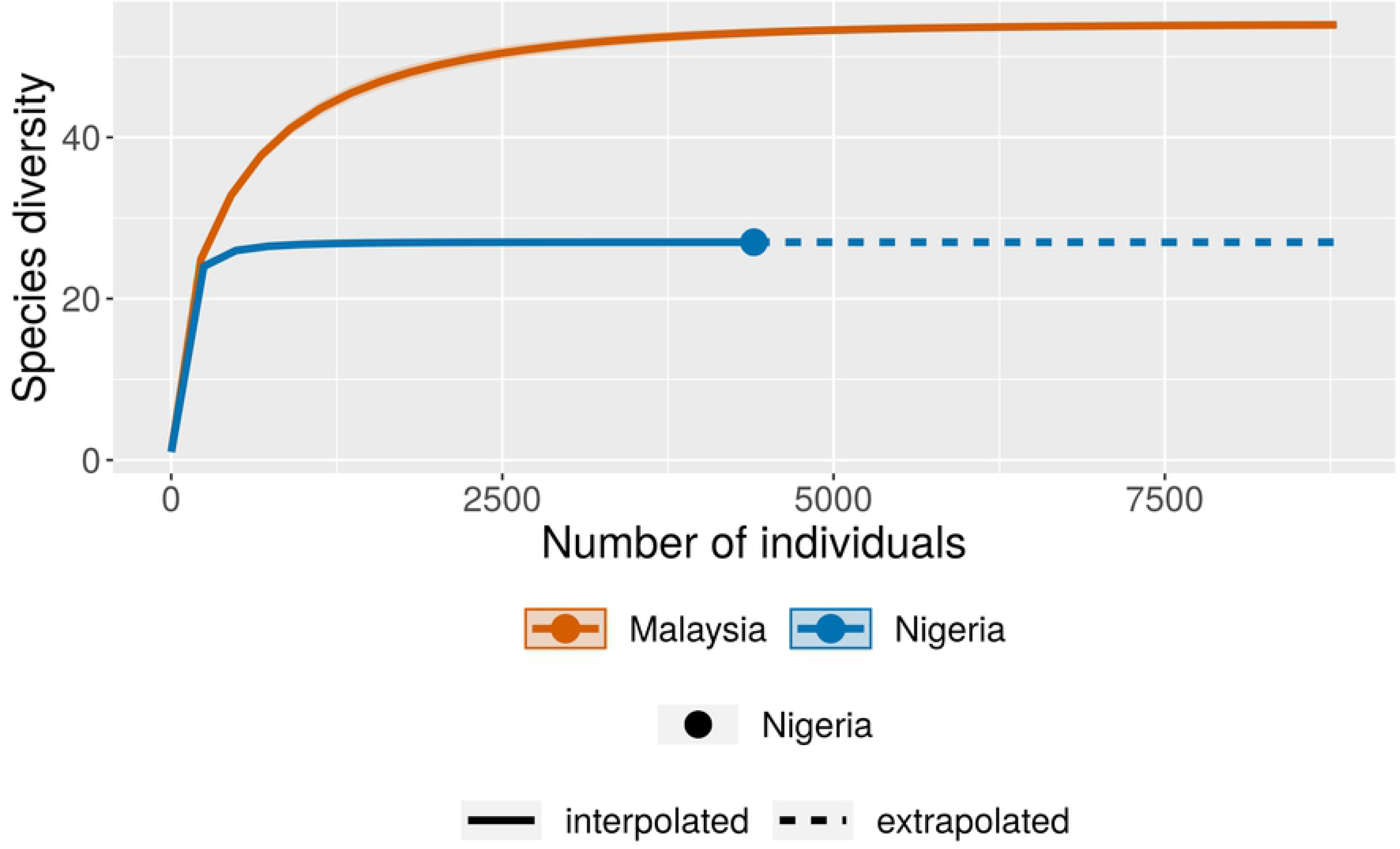
Individual-based rarefied-extrapolated species richness curves for the combined forests of both Malaysia and Nigeria.

**Table 1:**
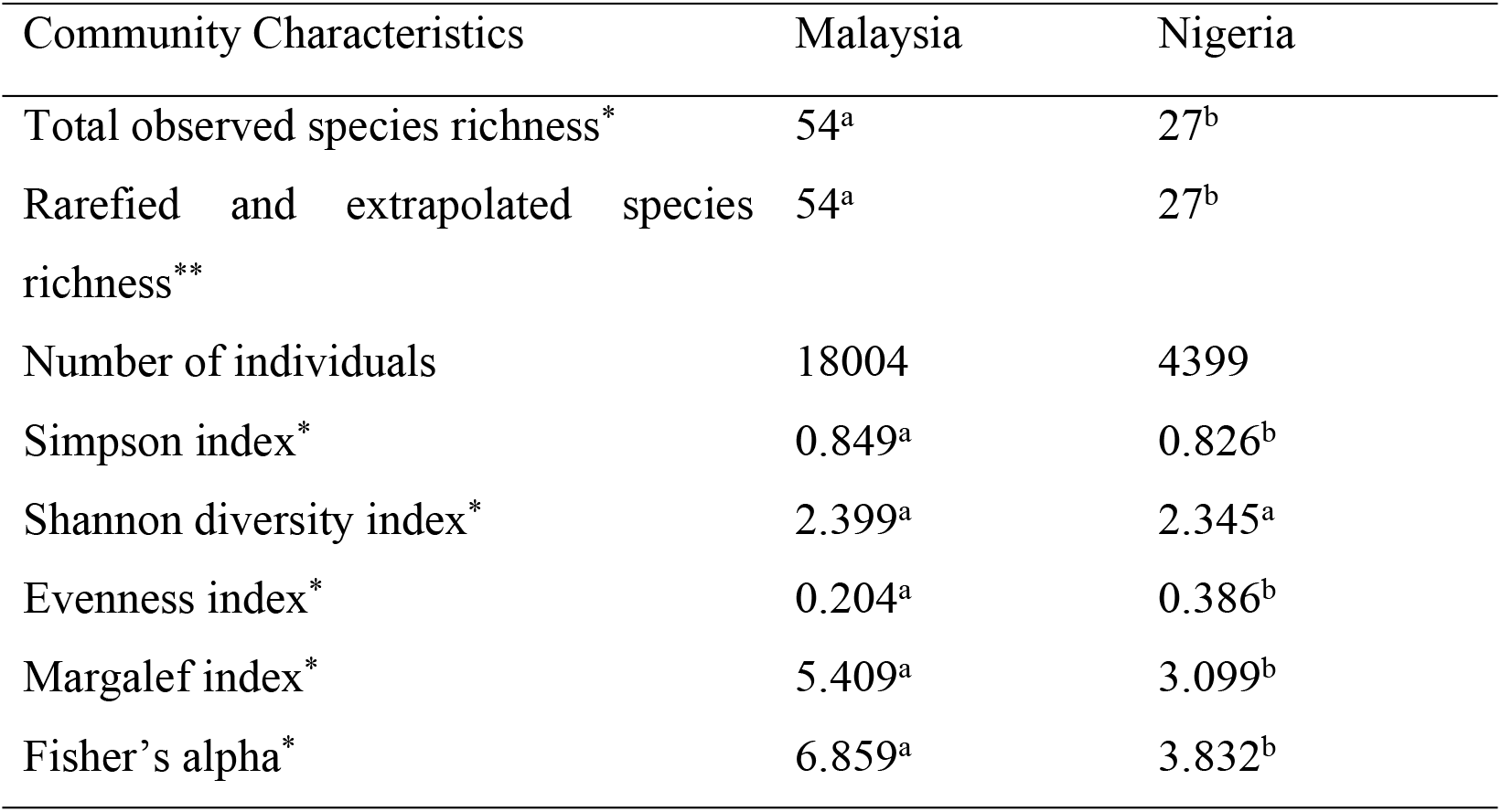
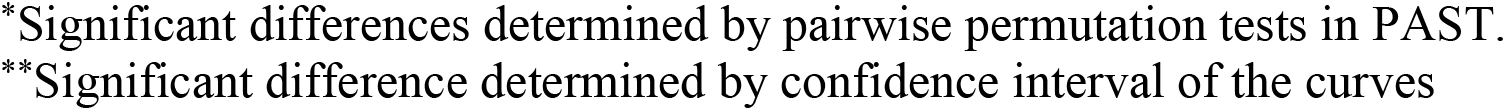
comparison of ferns community characteristics between Malaysia and Nigeria

| Community Characteristics | Malaysia | Nigeria |
| --- | --- | --- |
| Total observed species richness* | 54<sup>a</sup> | 27<sup>b</sup> |
| Rarefied and extrapolated species richness** | 54<sup>a</sup> | 27<sup>b</sup> |
| Number of individuals | 18004 | 4399 |
| Simpson index* | 0.849<sup>a</sup> | 0.826<sup>b</sup> |
| Shannon diversity index* | 2.399<sup>a</sup> | 2.345<sup>a</sup> |
| Evenness index* | 0.204<sup>a</sup> | 0.386<sup>b</sup> |
| Margalef index* | 5.409<sup>a</sup> | 3.099<sup>b</sup> |
| Fisher's alpha* | 6.859<sup>a</sup> | 3.832<sup>b</sup> |

\*Significant differences determined by pairwise permutation tests in PAST.
\*\*Significant difference determined by confidence interval of the curves

As for the individual forests in Malaysia, it was observed that the Bukit Hijau forest has a significantly higher number of ferns (44) than the USM campus forest (23). This trend was also observed in all the diversity indices of the two forests except for the evenness index which showed no significant difference between them (Table 2, Fig. 3b). In Nigeria, OAU campus forest was observed to have a significantly higher number of ferns (24) than Ikogosi warm spring forest (11). The same trend was also observed in all other diversity indices measured except the Simpson index (Table 2, Fig. 3c). These observed differences between the individual forests of each country could be due to other factors than climate. This is because the two study sites in each country are influenced by almost similar climatic conditions with slight differences. The most suspicious factor could be the extent of human disturbances in each forest [9]. For example, the Ikogosi warm spring forest is a recreational forest which receives so many tourists daily with no strict protection of the forest species. The reverse is the case for Bukit Hijau forest which has some measures of protection for its forest species [17].

**Figure 3b:**
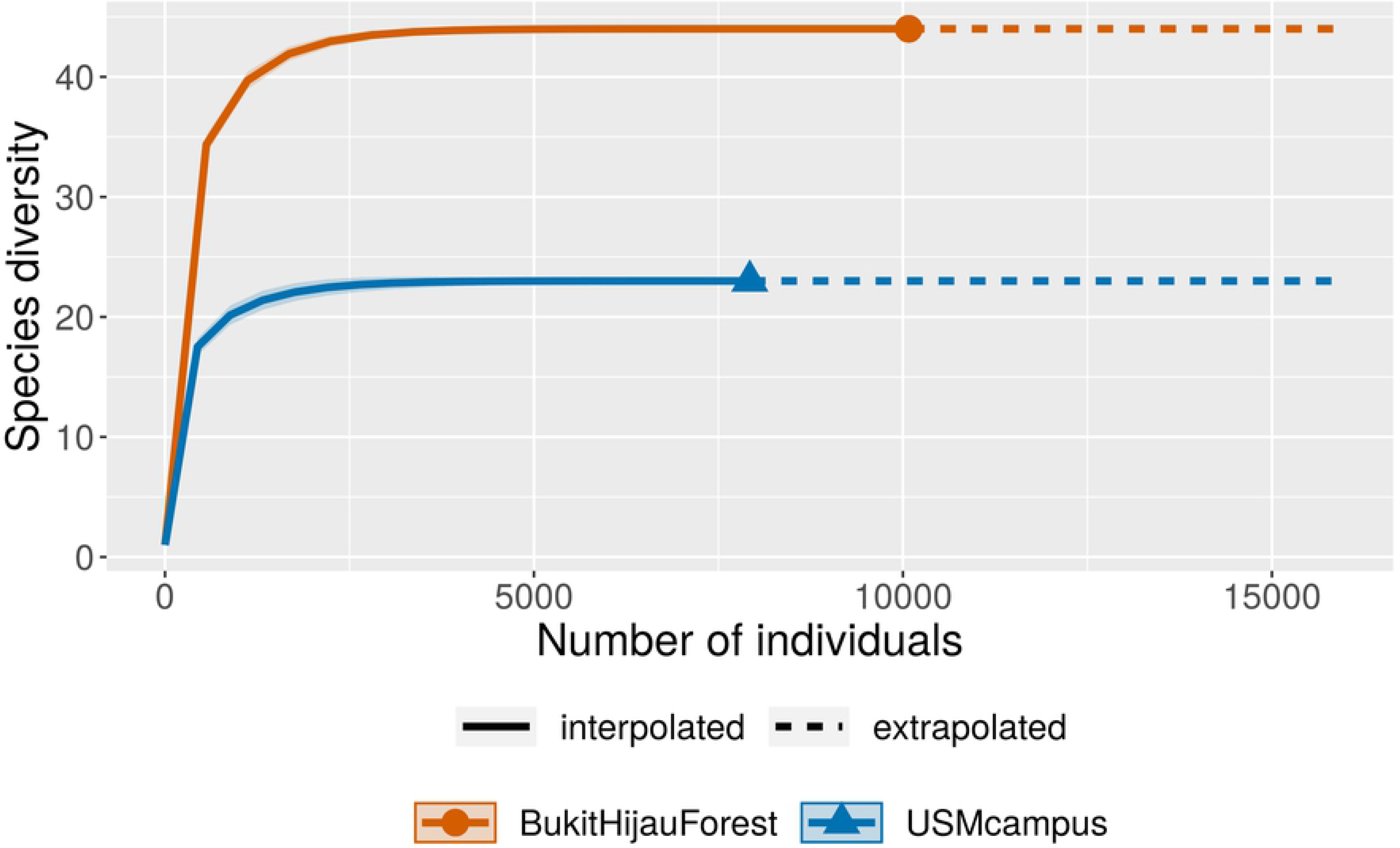
Individual-based rarefied-extrapolated species richness curves for the individual forests in Malaysia

**Figure 3c:**
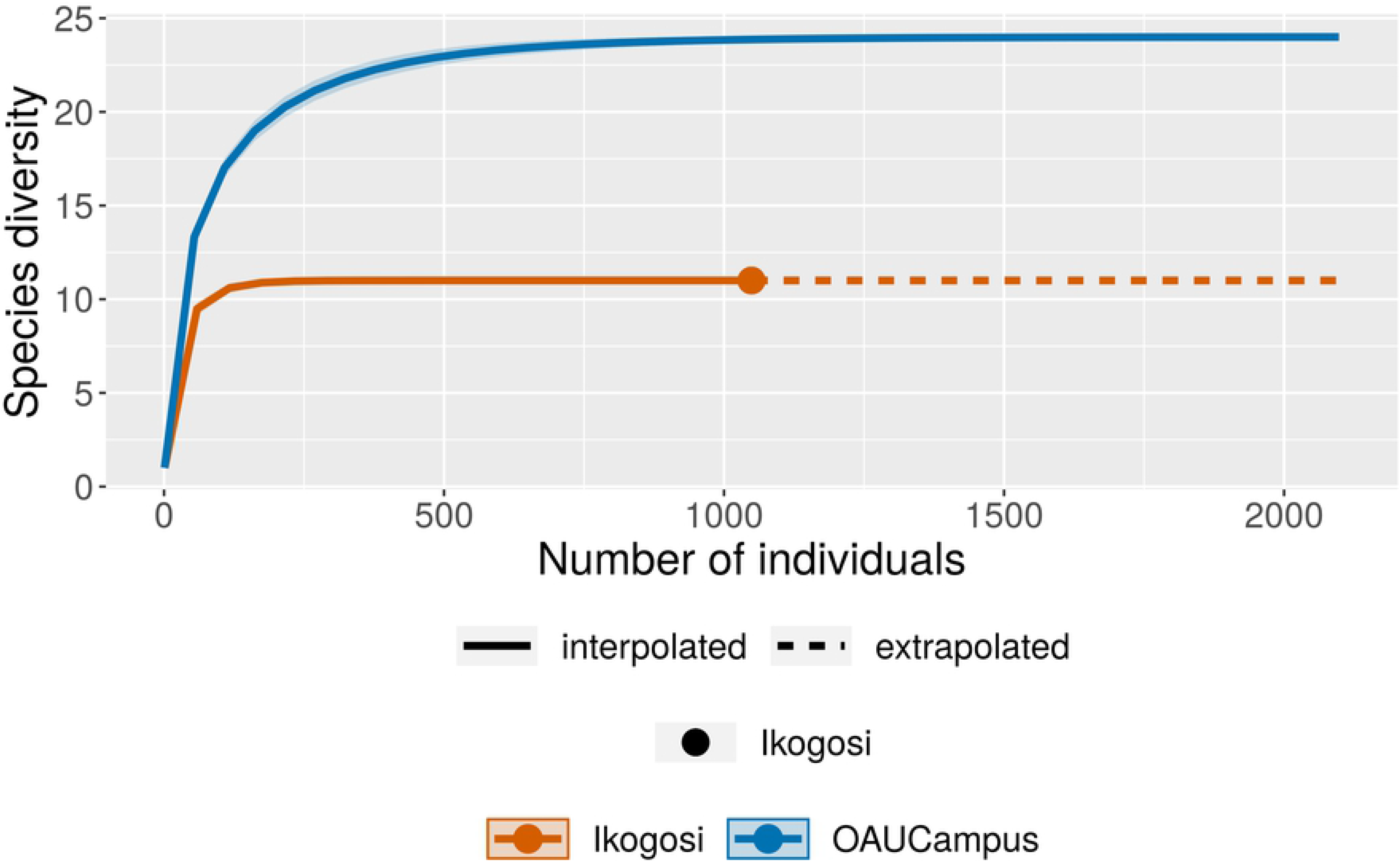
Individual-based rarefied-extrapolated species richness curves for the individual forests in Nigeria.

**Table 2:** comparison of ferns community characteristics between individual sampling sites in Malaysia and Nigeria

**Malaysia**
| Community Characteristics | USM Campus forest | Bukit Hijau forest |
| --- | --- | --- |
| Total observed species richness* | 23<sup>a</sup> | 44<sup>b</sup> |
| Rarefied and extrapolated species richness** | 23<sup>a</sup> | 44<sup>b</sup> |
| Number of individuals | 7925 | 10079 |
| Simpson index* | 0.762<sup>a</sup> | 0.874<sup>b</sup> |
| Shannon diversity index* | 1.828<sup>a</sup> | 2.517<sup>b</sup> |
| Evenness index* | 0.271<sup>a</sup> | 0.282<sup>a</sup> |
| Margalef index* | 2.450<sup>a</sup> | 4.665<sup>b</sup> |
| Fisher's alpha* | 2.907<sup>a</sup> | 5.913<sup>b</sup> |

**Nigeria**
| Community Characteristics | OAU Campus forest | Ikogosi warm spring forest |
| --- | --- | --- |
| Total observed species richness* | 24<sup>a</sup> | 11<sup>b</sup> |
| Rarefied and extrapolated species richness** | 24<sup>a</sup> | 11<sup>b</sup> |
| Number of individuals | 3327 | 1049 |
| Simpson index* | 0.753<sup>a</sup> | 0.778<sup>a</sup> |
| Shannon diversity index* | 2.100<sup>a</sup> | 1.829<sup>b</sup> |
| Evenness index* | 0.340<sup>a</sup> | 0.566<sup>b</sup> |
| Margalef index* | 2.836<sup>a</sup> | 1.438<sup>b</sup> |
| Fisher's alpha* | 3.499<sup>a</sup> | 1.714<sup>b</sup> |

\*Significant differences determined by pairwise permutation tests in PAST.
\*\*Significant difference determined by confidence interval of the curves

Generally, terrestrial ferns are the dominant ferns in Malaysian and Nigerian forests having 37 and 17 species respectively while aquatic ferns are the least (Table 3). The dominant nature of the terrestrial ferns could also be an indicator to the lesser degree of disturbance of the forests in the two countries when considered on a larger scale. Researchers have confirmed the dominant nature of terrestrial ferns in less-disturbed forests [15, 17, 25, 26]. The non-metric multidimensional scaling analysis showed that the Malaysian and Nigerian forests are more associated with terrestrial and epiphytic ferns than the other life forms of ferns (Fig. 4).

**Figure 4:**
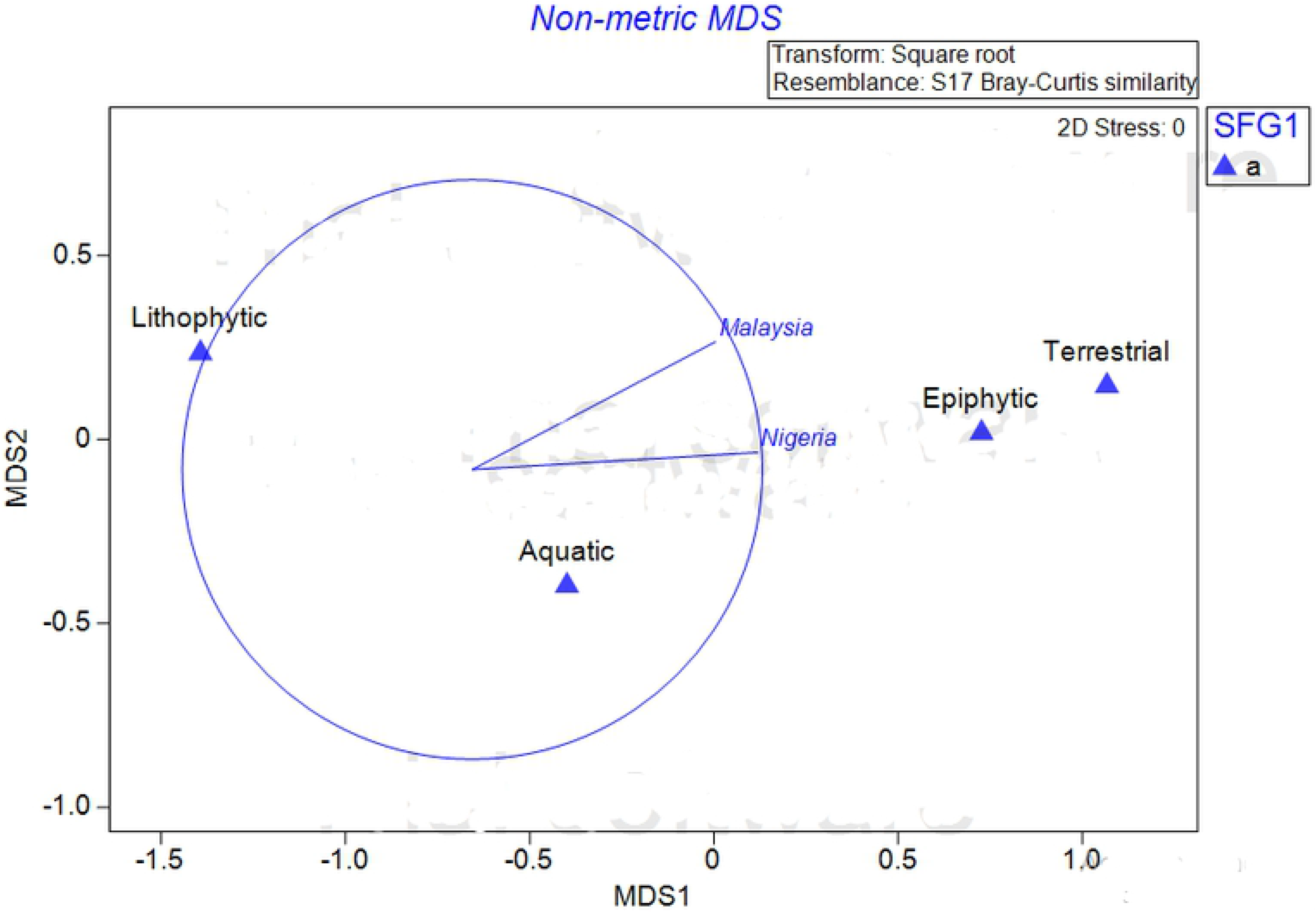
nMDS showing the association between the ferns life forms and the two countries.

**Table 3:**
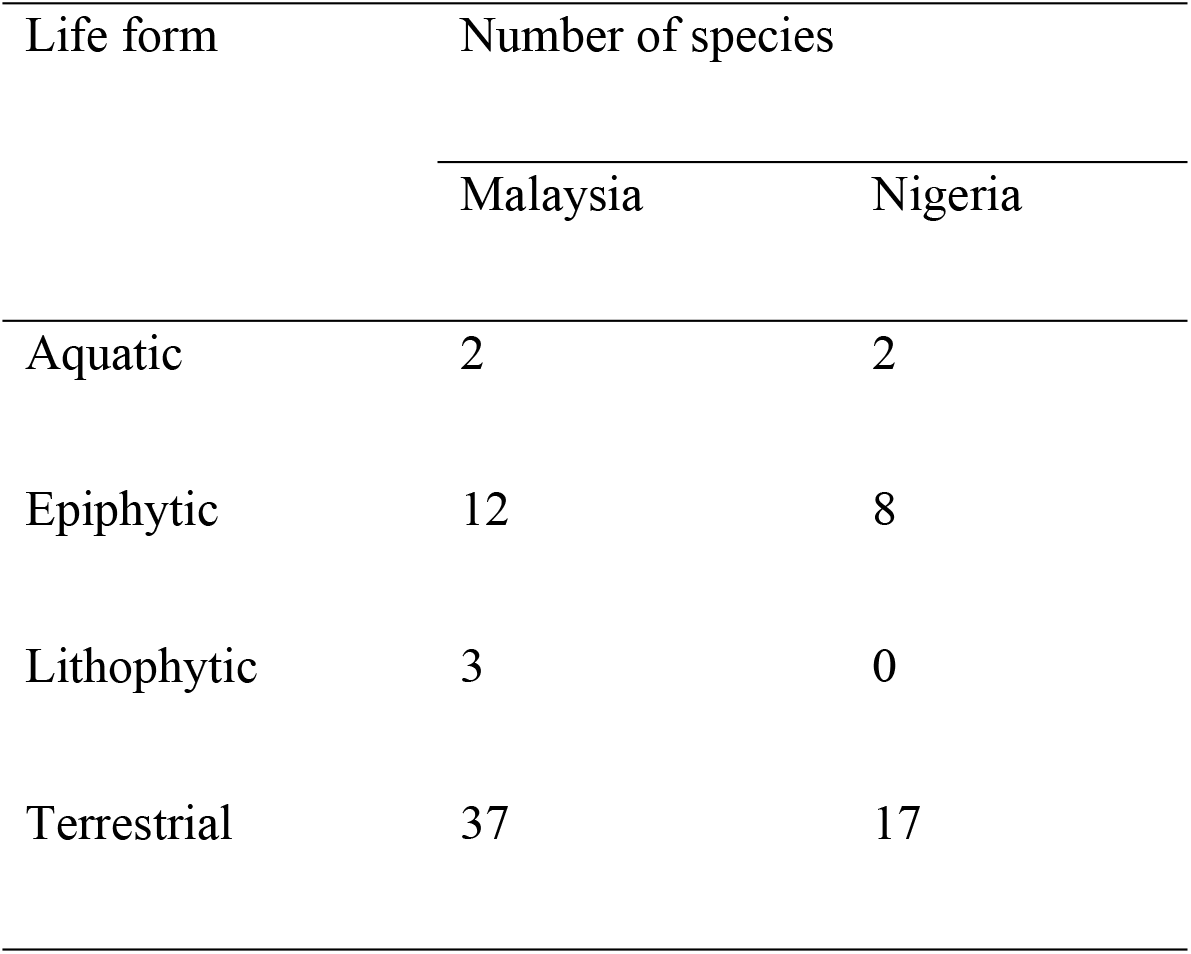
Life forms of the ferns in Malaysia and Nigeria

It should be noted that some of these identified ferns exhibited more than one life form. For example, *Pteris vittata* existed as both terrestrial and lithophytic fern while *Nephrolepis biserrata* existed as both terrestrial and epiphytic fern in Malaysia and Nigeria. Some ferns have also been reported to have the ability to exist in different life forms [27]. The PCA showed that the principal components (PC) 1 and 2 contributed to 93.52% and 6.48% of the total variations respectively (Fig. 5). However, Malaysian forests contributed more significantly to the PC 1 and Nigerian forests contributed to PC 2. This invariably shows that Malaysian forests contributed largely to the overall variations in the fern species. *Pyrrosia lanceolata* and *Drynaria quercifolia* were observed to be the ferns with the highest relative frequencies of 25.36% and 24.59% respectively in the combined forests of Malaysia (Table 4). In Nigeria, *Pneumatopteris afra* with the relative frequency of 35.62% was observed to be the highest.

**Figure 5:**
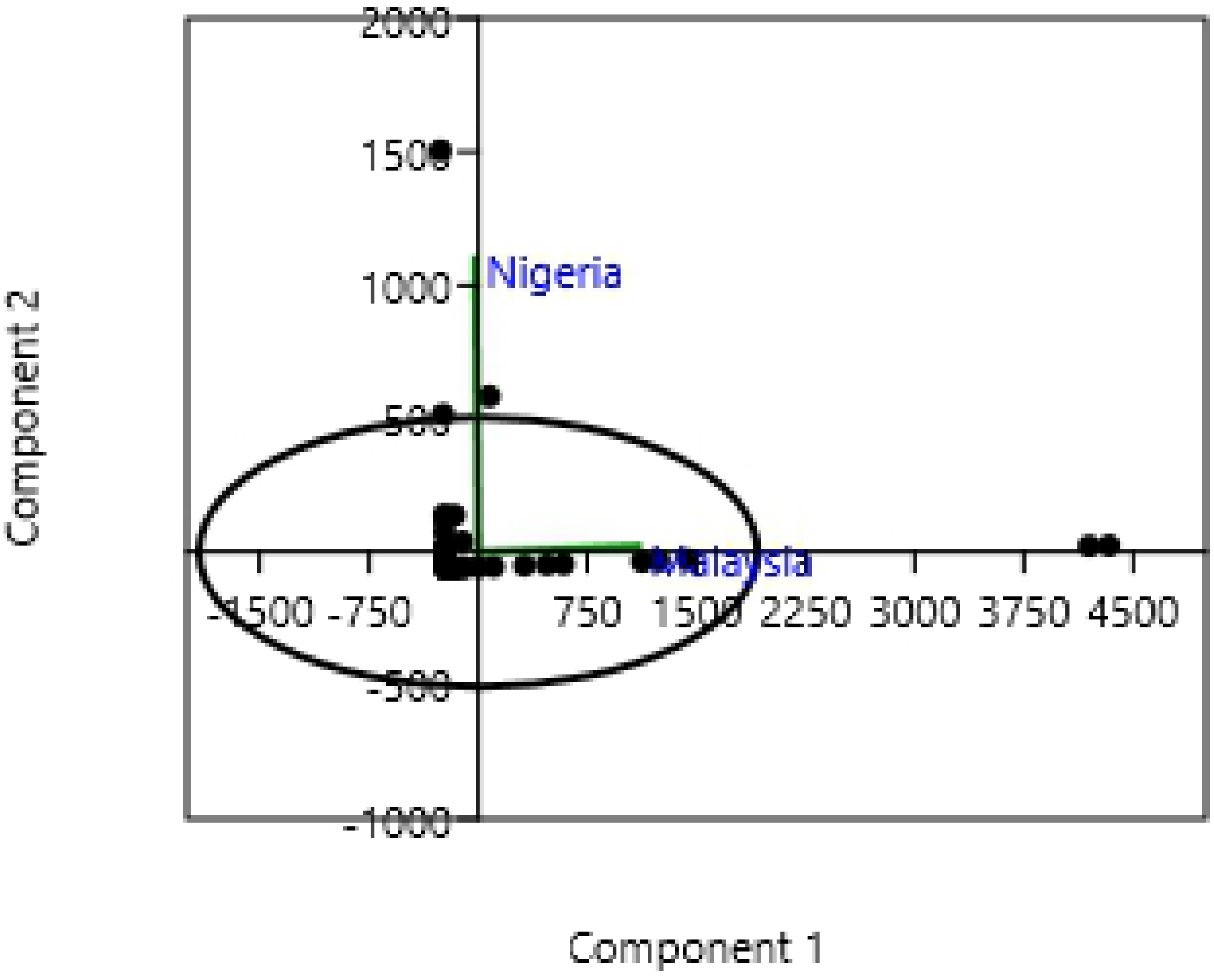
Principal component analysis showing the variations in the distribution of fern species in the two countries.

**Table 4:**
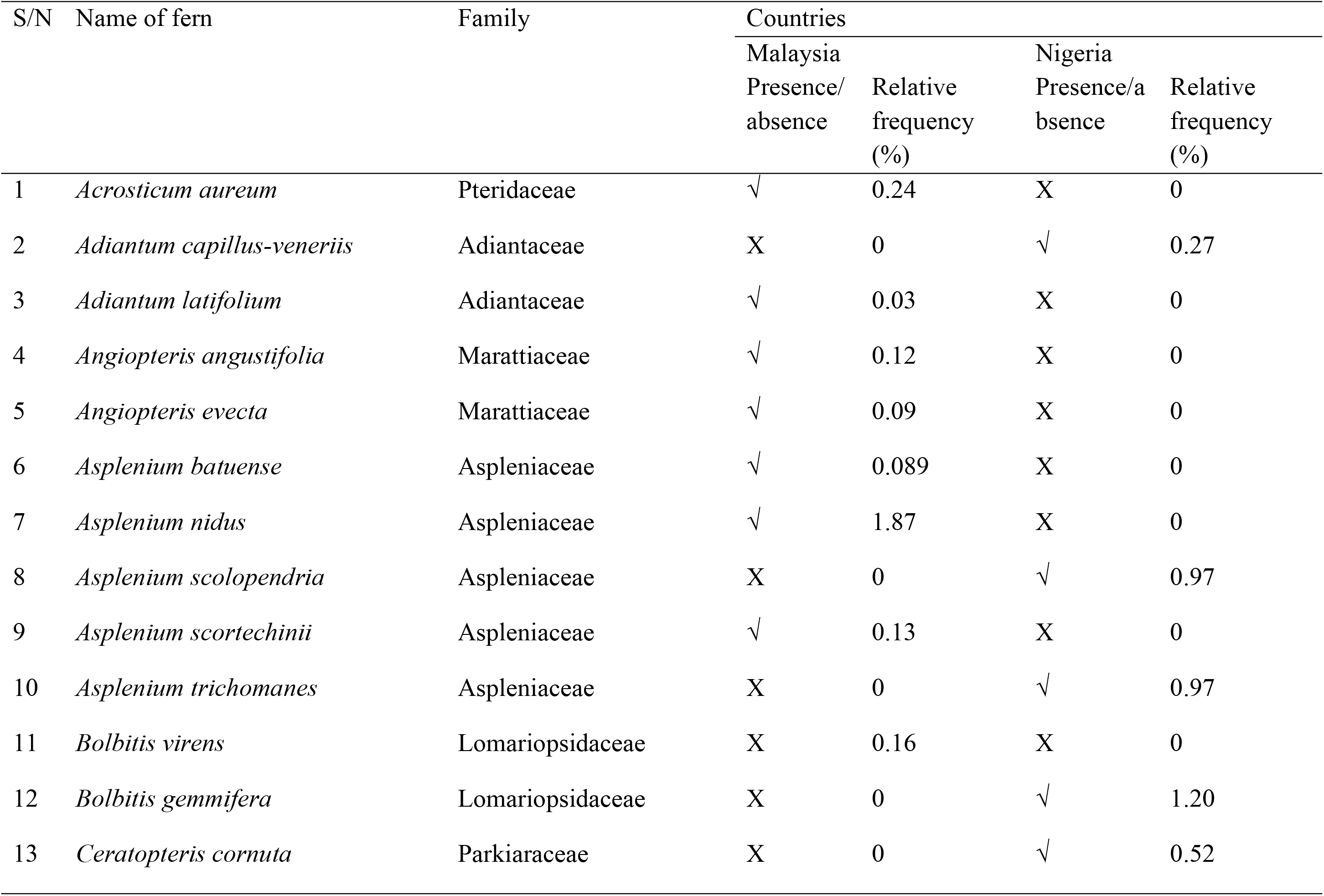
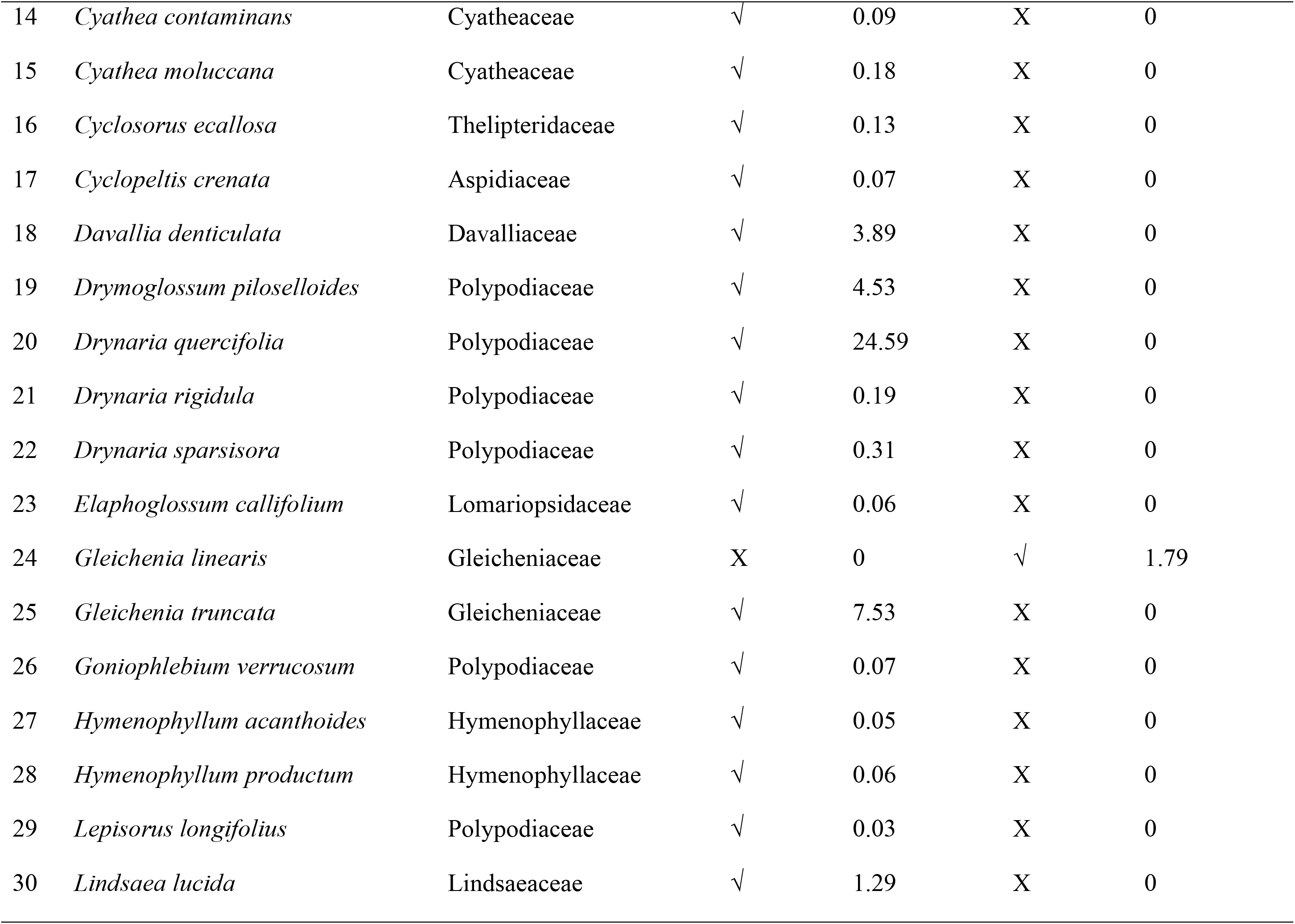
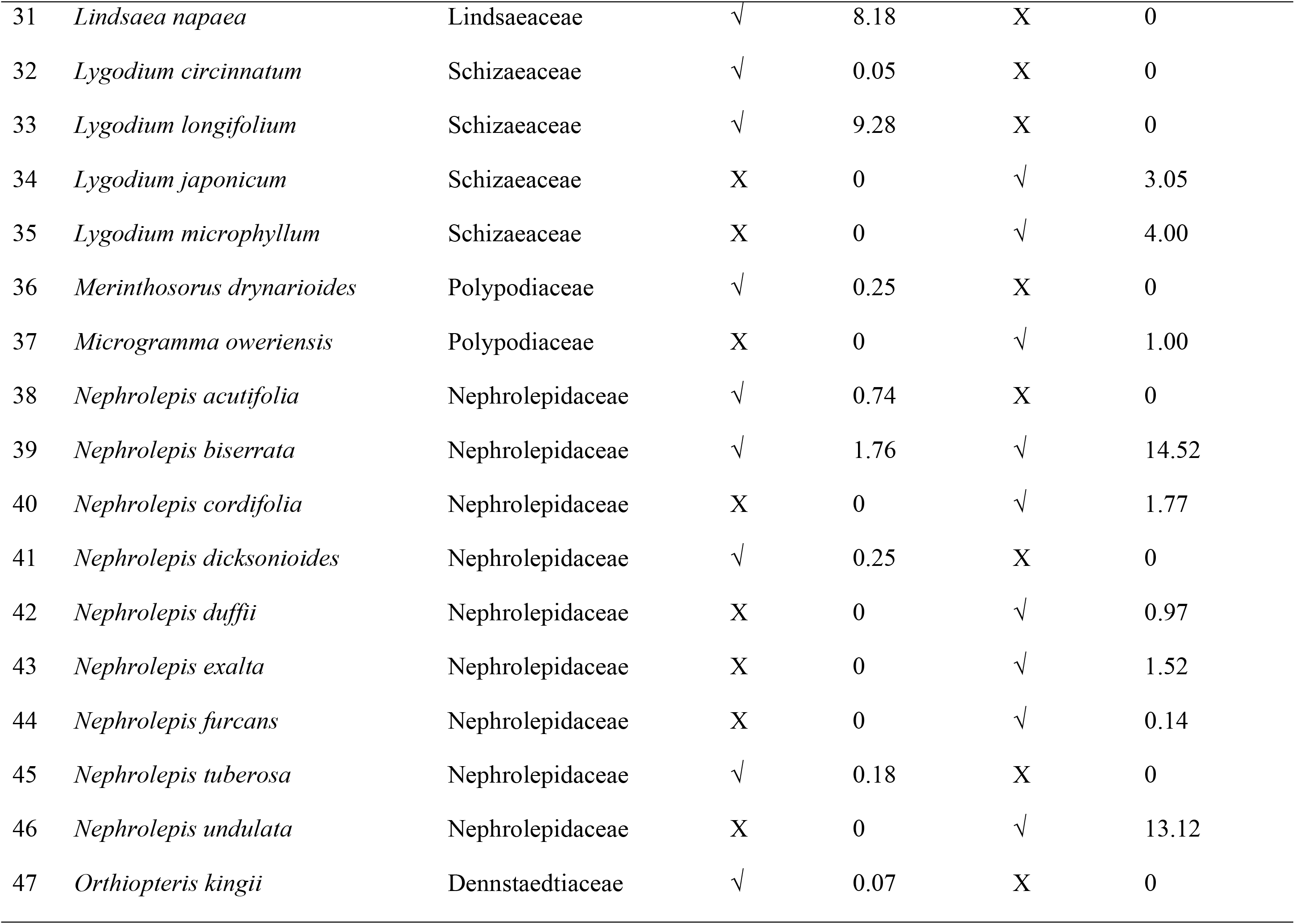
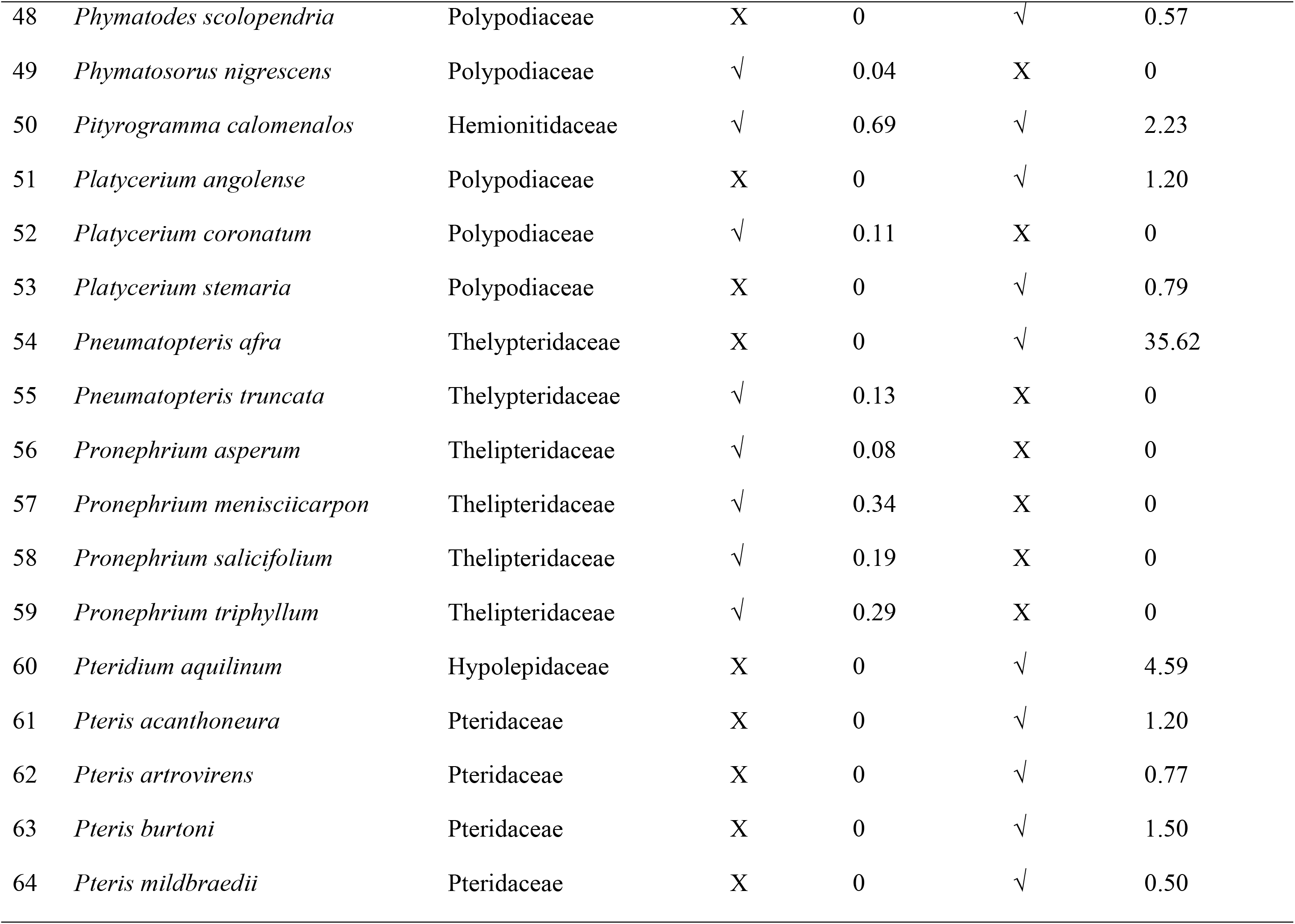
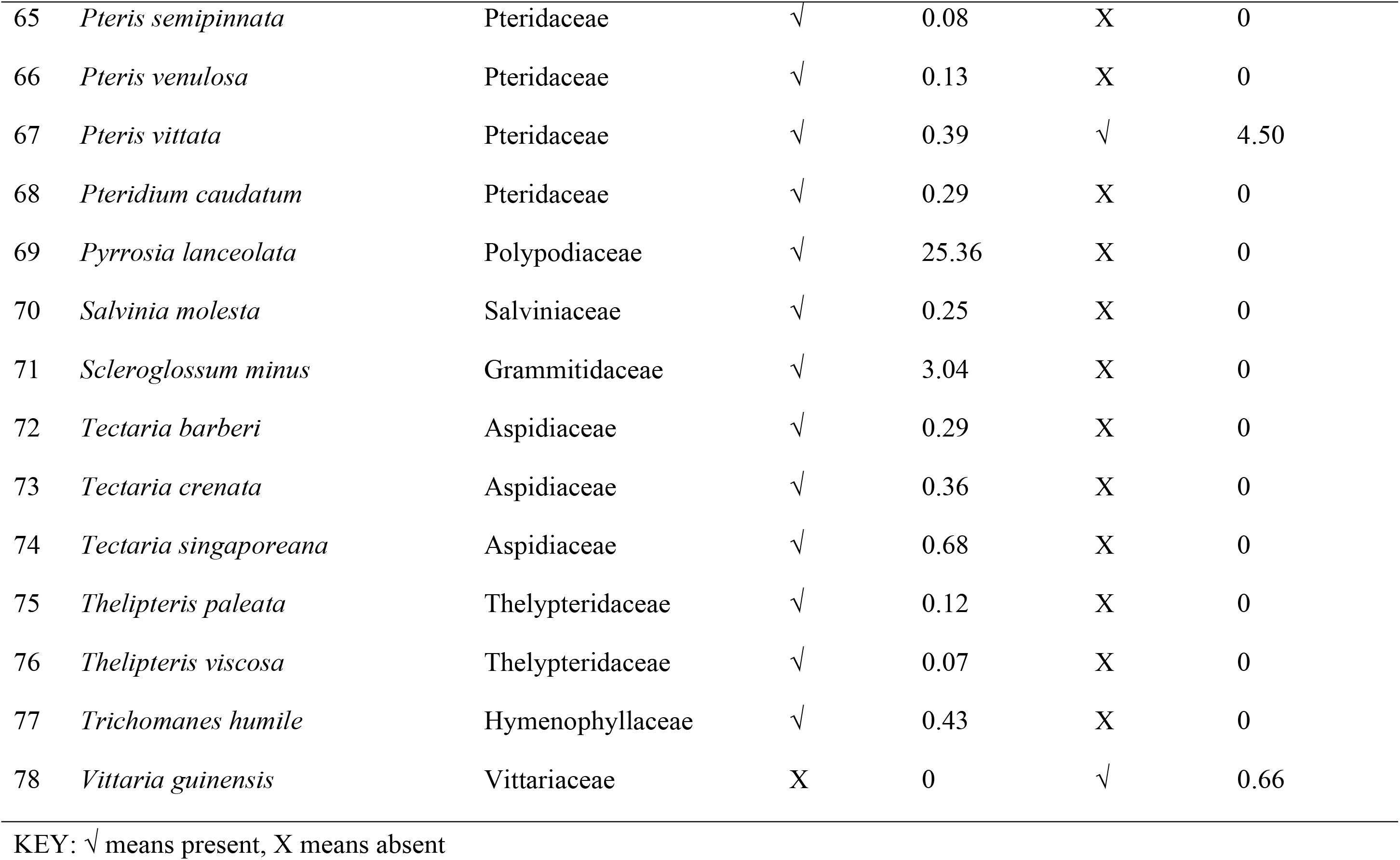
The distribution of ferns observed in the study areas

The common fern species shared by the two countries include *Nephrolepis biserrata, Pityrogramma calomelanos,* and *Pteris vittata.* This makes the Sorenson similarity index of the fern species between the two countries to be 7.41%. This is a very low index of similarity which indicates that the two countries, though within the tropics, do not share many fern species in common. This may be due to differences in their evolutionary or historical processes, plus climatic factors of the two regions [28]. However, the similarity in the ferns composition between the two countries increased as the taxonomic level increased (species: 7.41%, genus: 12.77%, family: 70.96%). The family Polypodiaceae represented by seven genera and nine species is the dominant family of ferns in Malaysia whereas Nephrolepidaceae represented by one genus and six species is the dominant family in Nigeria (Table 5). The dominance of the family Polypodiaceae in Peninsular Malaysia has also been reported by previous studies [17, 7].

**Table 5:**
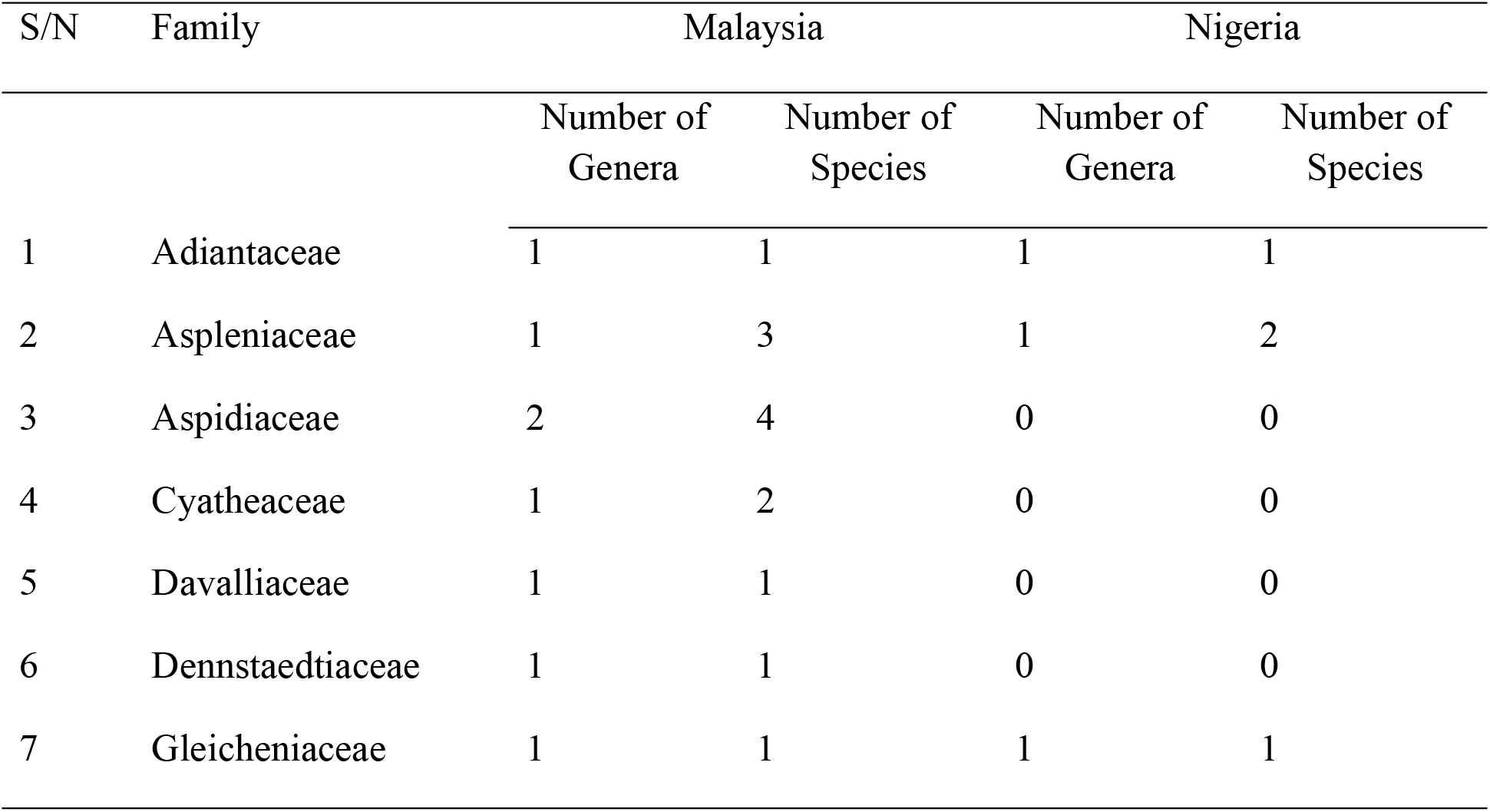
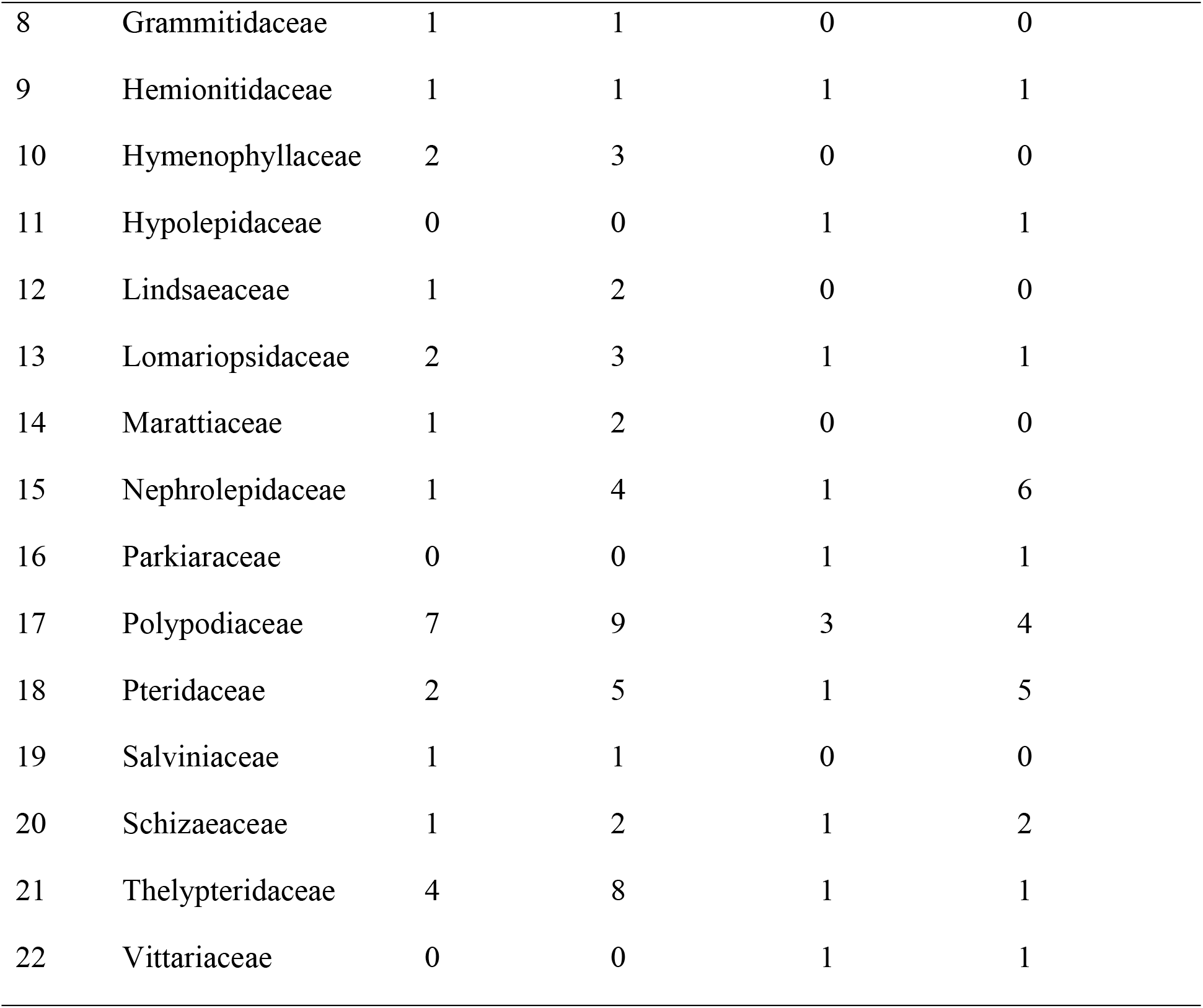
Checklist of families with respective numbers of general and species

## Conclusion

The results of this study indicated that the Malaysian forests provide favourable microclimatic conditions which supported significantly higher ferns diversity and richness than Nigerian forests. There are distinct differences in the ferns community composition between the two countries which led to a very low similarity index at the species and genus levels. These observed differences in the species composition may be explained by differences in climatic and edaphic factors, as well as historical, and evolutionary processes between the two biogeographic areas. We thereby recommend that future researches should include forests of other countries to have a robust comparison of variations in ferns community assemblages between Asia and Africa.

## Acknowledgement

The authors hereby acknowledge the Universiti Sains Malaysia for creating enabling environment and for the financial support towards the success of this research.

## Notes

### Competing Interest Statement

The authors have declared no competing interest.

